# Structure-Based Drug Design of Novel Inhibitors Targeting the Thiamine Biosynthesis Pathway in Bacteria

**DOI:** 10.64898/2026.10.05.756923

**Authors:** Nikil Krishna, Venkatasubramanian Ulaganathan

## Abstract

Current generation antibiotics are limited by poor target selectivity and are often mismanaged, leading to widespread antimicrobial resistance. The bacterial thiamine biosynthesis pathway offers compelling alternative since it is highly essential for bacterial survival and yet absent in humans. Genetic perturbations within pathway lead to documented auxotrophy and virulence defects, indicating low mutational tolerance within this pathway. Thiamine phosphate synthase is a key bottleneck enzyme performing the crucial role of coupling thiazole and pyrimidine moieties to form thiamine monophosphate. To develop a suitable inhibitor against it, a structure-based drug design pipeline was implemented. High-throughput virtual screening of nearly 1.5 million molecules provided sufficient exploration of chemical space. Further, R-group substitution was utilized for lead optimization. Structure-Activity Relationship analysis upon molecular mutants reveals patterns among R-group interactions to catalytic residues across various scaffolds. Additionally, ADMET screening and validation using molecular dynamics simulations resulted in a potent orally bioavailable broad-spectrum bactericidal molecule.

## 1. INTRODUCTION

According to WHO, antimicrobial resistant infections cause 700,000 deaths annually and is expected to reach 10,000,000 deaths per annum by 2050 if left unchecked^1^. The primary drivers are over-exploitation of antibiotic usage focused on a limited set of targets within the bacterial life cycle, often the central dogma and the cell wall^2,3^. Further drug discovery focusing on these targets is futile and hence additional exploration of druggable targets, particularly within metabolic pathways, is essential to overcome the current pace of resistance development^4^.

### 1.1 The Thiamine (Vitamin B1) Molecule

Thiamine is an essential cofactor for multiple enzymatic systems including but not limited to, pyruvate dehydrogenase complex within TCA cycle, transketolase within pentose phosphate pathway, and alpha-ketoglutarate dehydrogenase within amino acid biosynthesis^1,5^. Vertebrates, including humans, exclusively obtain thiamine from dietary sources and lack all the enzymatic machinery required for denovo synthesis^1,6^. This divergence creates an opportunity for the development of highly selective antimicrobial agents exhibiting minimal human toxicity. Recent comprehensive phylogenetic analysis of 154,000 bacterial genomes revealed that thiamine biosynthetic genes are widely distributed across bacteria and present in all clinically relevant pathogens^7^. Further, prior studies have confirmed the high druggability of key enzymes within the pathway^8,9^.

### 1.2 The Thiamine Biosynthesis Pathway and Key Enzymatic Targets

Thiamine biosynthesis pathway involves the parallel synthesis of two heterocyclic compounds: thiazole and pyrimidine, which are coupled to form thiamine monophosphate, which is then phosphorylated to generate thiamine diphosphate^9,10^ Three bottleneck rate-limiting steps within the pathway are Dxs (1-deoxy-D-xylulose-5-phosphate synthase) coupling pyruvate and glyceraldehyde-3-phosphate to form deoxy-D-xylulose 5-phosphate, ThiE (Thiamine phosphate synthase) coupling thiazole and pyrimidine to form thiamine phosphate and ThiL (Thiamine phosphate kinase) phosphorylating thiamine monophosphate to generate thiamine diphosphate^1^ (Figure 1). Further, genetic and biochemical studies show that mutations of any kind within these enzymes result in growth defects and reduced virulence within *in vivo* animal models^5,8^.

**Figure 1:**
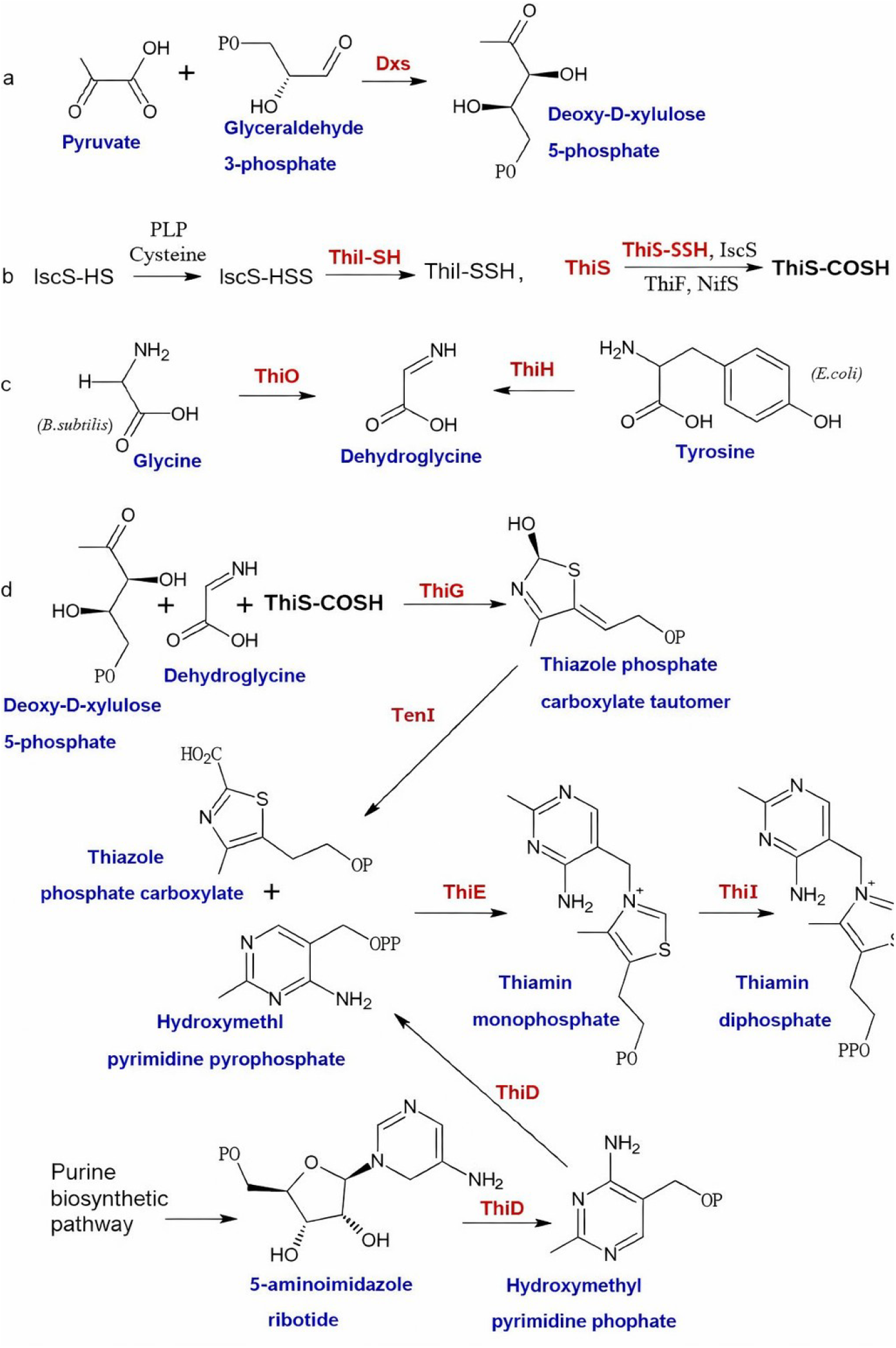
Thiamine Biosynthesis Pathway

### 1.3 Selection of ThiE as the Primary Target for Investigation

ThiE catalyzes a crucial coupling reaction of thiazole and pyrimidine precursors to form the heterocyclic thiamine phosphate. This step cannot be circumvented through other metabolic pathways and hence, limiting the potential for drug resistance by metabolic re-routing. Prior biochemical screening studies have explored the druggability of this enzyme and had successfully identified thiazolidine-containing inhibitors of *Mycobacterium tuberculosis* ThiE with low micromolar potency. Multiple sequence alignment and structural superposition analysis further reveal identical evolutionary conservation of active site residues among ThiE orthologs, suggesting that inhibitors designed against ThiE may possess broad-spectrum activity. Furthermore, ThiE and its homologs are absent in human and mammalian hosts. Due to these considerations, ThiE was selected as the reference target for this workflow with *E. coli* serving as the model system due to its availability of detailed genomic and proteomic resources ^10– 12^.

## 2. MATERIALS AND METHOD

### 2.1 Protein Target Characterization and Structural Analysis

Multiple sequence alignment using mBED algorithm implemented at the Clustal Omega web server (https://www.ebi.ac.uk/jdispatcher/msa/clustalo) and structural superposition using PyMOL (version 2.5.4) was performed on ThiE orthologs from three distinct phylogenetic lineages and clinical importance, including *Escherichia coli K12* (UniProt ID P30137), *Mycobacterium tuberculosis* (UniProt ID P9WG75), and *Bacillus subtilis* (UniProt ID P39594), was performed to identify active site conservation. Root mean squared deviation was computed over aligned protein backbone^13^ (Supplemental Figure 1).

### 2.2 Chemical Database Preprocessing and Drug-Likeness Filtering

Before initiating virtual screening, database pre-processing and filtering was performed to reduce effective chemical space focusing on the subset of compounds with favorable pharmacokinetic properties to proceed to docking^14,15^. Compound satisfying drug-likeliness criteria exhibit improved attrition rates and higher probability of oral bioavailability^14,16^. All the required descriptors were calculated using RDKit library. The chemical compound library was obtained from ChEMBL (version 36, https://www.ebi.ac.uk/chembl/), comprising of experimentally validated bioactive compounds with favorable chemical diversity metrics and synthetic accessibility. Initially comprising of 2,783,916 compounds, two complementary drug-likeliness systems were sequentially applied to identify compounds with high predicted favorable oral bioavailability.

#### Lipinski’s Rule of Five Filtering

Lipinski’s Rule of Five, formulated by Christopher Lipinski in 1997, constitutes the most widely employed heuristic for predicting oral drug-likeness in early-stage drug discovery^17,18^. In accordance with this framework, the present investigation applied the following thresholds, molecular weight (0-500 Da), LogP (0.4-5.6), hydrogen bond donors (0-5), and hydrogen bond acceptors (0-10).

#### Veber’s Rules Filtering

Veber’s Rules represent a complementary set of drug-likeness criteria derived from analysis of over 1,100 drug candidates evaluated for oral bioavailability at pharmaceutical organizations^18^. Veber’s analysis demonstrated that reduced molecular flexibilitys (measured by the number of rotatable bonds) and low topological polar surface area (TPSA) or total hydrogen bond count represent important independent predictors of good oral bioavailability, particularly for compounds with molecular weight < 400 Da. Applied Veber’s criteria were: rotatable bonds ≤ 10 and topological polar surface area ≤ 140 Å^2^, which define regions of chemical space exhibiting high probability of favorable oral bioavailability. Recent comprehensive analysis of FDA-approved drugs has confirmed that Veber’s Rules exhibit superior predictive power for oral bioavailability compared to Lipinski’s Rule of Five alone, with approximately 85% of marketed oral drugs conforming to Veber’s criteria compared to 66% conforming to strict Lipinski’s Rule of Five^19^. The incorporation of both filtering systems provides orthogonal assessment of drug-likeness from complementary perspectives.

### 2.3 High-Throughput Virtual Screening Using GPU-Accelerated Molecular Docking

Virtual screening through computational molecular docking enables efficient prioritization and ranking of compounds from large chemical libraries based on predicted binding affinity to target protein^22-24^. High-throughput GPU-accelerated molecular docking was performed using Unidock^25^ utilizing Autodock Vina^26^ scoring function. Docking calculations employed a rigid body docking algorithm^22,23^ (exhaustiveness 16) utilizing a docking grid box constructed using coordinates 1.0686, 8.6604, 0.1427 and dimensions 30 Å, 30 Å, 30 Å. The molecules were sorted according to their binding affinities and top 10 ranking molecules were used for further lead optimization.

### 2.4 Structure-Activity Relationship Analysis and R-Group Optimization

To understand how atomic changes to the ligand affects the calculated binding affinity of the complex, R-group substitution and structure-activity relationship analysis was utilized to identify which chemical substituents and functional groups within the ligand enhance binding affinity, enabling targeted modification of lead structures to achieve superior potency^27,28^. Top 10 ranking scaffolds from the virtual screening stage were selected as lead compounds. Within each structure, two growth sites, R1 and R2, were selected as scaffold exit vectors for 10 distinct R-group substitutions listed as follows: methyl (hydrophobic), tert-butyl (hydrophobic), fluoro (electrophilic), chloro (electrophilic), hydroxyl (hydrophilic), methoxy (hydrophilic), carboxylic acid (acidic), amino (basic), carboxamide (hydrophilic), and pyridyl (aromatic and hydrophobic). These substituents were selected to cover comprehensive chemical diversity. Next, to accurately predict the difference in biological activity for the molecular mutants, Boltz-2, a state-of-the-art biomolecular foundation transformer model was utilized. The 1,000 molecular mutants were co-folded with the protein via induced fit using the PairFormer module and their biological activity were predicted using the affinity module. For each compound, the effects of the substitution patterns on predicted binding affinity were analyzed.

### 2.5 Comprehensive ADMET Profiling Using Multi-Criteria Decision Analysis

Top 20 ranking lead molecules were selected for further ADMET screening. ADMET (absorption, distribution, metabolism, excretion, toxicity) screening is performed to identify and eliminate compounds with poor pharmacokinetic properties and toxicity^29,30^. Comprehensive ADMET profiling of top 20 lead compounds using ADMETlab 3.0 web server, a machine learning-based platform employing directed message passing neural network (DMPNN) for accurate prediction of nearly 90 distinct ADMET endpoints, including but not limited to molecular descriptors (molecular weight, logP, topological polar surface area, number of hydrogen bond donors and acceptors, number of rotatable bonds, number of aromatic rings, number of heavy atoms), ADMET properties (permeability, blocking tendency, penetration likelihood of blood-brain barrier, hepatic clearance, renal excretion), Toxicity prediction (hepatotoxicity, acute toxicity, carcinogenicity, mutagenicity) and Medicinal rule compliance (Lipinski, Veber, and Ghose compliance)^31^.

In order to rank and sort the molecules given the high dimensionality of the ADMET output data, TOPSIS (Technique for Order of Preference by Similarity to Ideal Solution) was used. TOPSIS is a well-established multi-criteria decision analysis method that compares data based on their geometric closeness to an ideal solution. TOPSIS score based on the Euclidean distances between each compound and its ideal positive solution were calculated using a decision matrix containing the normalized ADMET values for each compound. The weighing schemes for TOPSIS reflecting the relative importance of these criteria included absorption and distribution properties (weightage 0.3), metabolism and excretion properties (weightage 0.25), toxicity assessments (weightage 0.35) and medicinal chemistry compatibility (weightage 0.1)^33^.

### 2.6 Molecular Dynamics Simulations for Binding Stability Assessment

Static docking conformations and calculations do not account for protein flexibility, entropic effects, and thermal fluctuations. Molecular dynamics simulations provide validation for binding pose stability under real physiological conditions. Hence, molecular dynamics simulations were performed on the protein-ligand complex of the selected molecule using OpenMM (version 8.0.1). The simulation parameters were set as follows:

The Amber19 and GAFF2 forcefield was used for protein and ligand parameterization, respectively. Explicit aqueous solvation was performed using TIP3P water model and 0.15 M counterions (Na^+^ and Cl^−^) were added to maintain electrostatic neutrality of the system in a rectangular grid box. The system was subjected to a two-stage equilibration protocol prior to production dynamics including NVT (constant number, volume, temperature) equilibration for 10 nanoseconds at 310 Kelvin with harmonic position restraints applied to protein and ligand heavy atoms with a force constant of 500 kJ/mol/nm^2^ and NPT (constant number, pressure, temperature) equilibration for 10 ns at 310 K and 1 bar pressure, applying positional restraints only to protein backbone atoms, allowing the system volume to equilibrate to appropriate density. Following equilibration, 100 ns of unrestrained molecular dynamics was performed in the NPT ensemble at 310 K and 1 bar, integrating Newton’s equations of motion using the velocity Verlet integrator with integration time step of 2 femtoseconds. Periodic boundary conditions were maintained throughout the simulations. Trajectory frames were collected every 10 picoseconds, generating 10,000 frames spanning the full 100 ns production run. Temperature and pressure were maintained constant using Langevin thermostat with a collision frequency of 1 ps^−^ and barostat with a pressure coupling constant of 1 ps. The trajectory was unwrapped and analysis was performed using MDAnalysis (version 2.5.0). Various stability metrics such as protein backbone root mean squared deviation, ligand all-atom root mean squared deviation, radius of gyration, and residue-level root mean squared fluctuation were calculated ^34-42^.

## 3. RESULTS AND DISCUSSION

### 3.1 Database Preprocessing

The sequential filtering process using Lipinski’s and Veber’s rules eliminated 1,360,102 compounds (48.9% of the original library) not satisfying the criteria and retaining 1,423,814 compounds for downstream operations (Figure 2). The retained compound subset exhibited improved molecular descriptors, such as mean molecular weight decreased from 695 to 358 Dalton, mean logP reduced to 2.15, and mean topological polar surface area decreased from 196 to 109 Å^2^.

**Figure 2:**
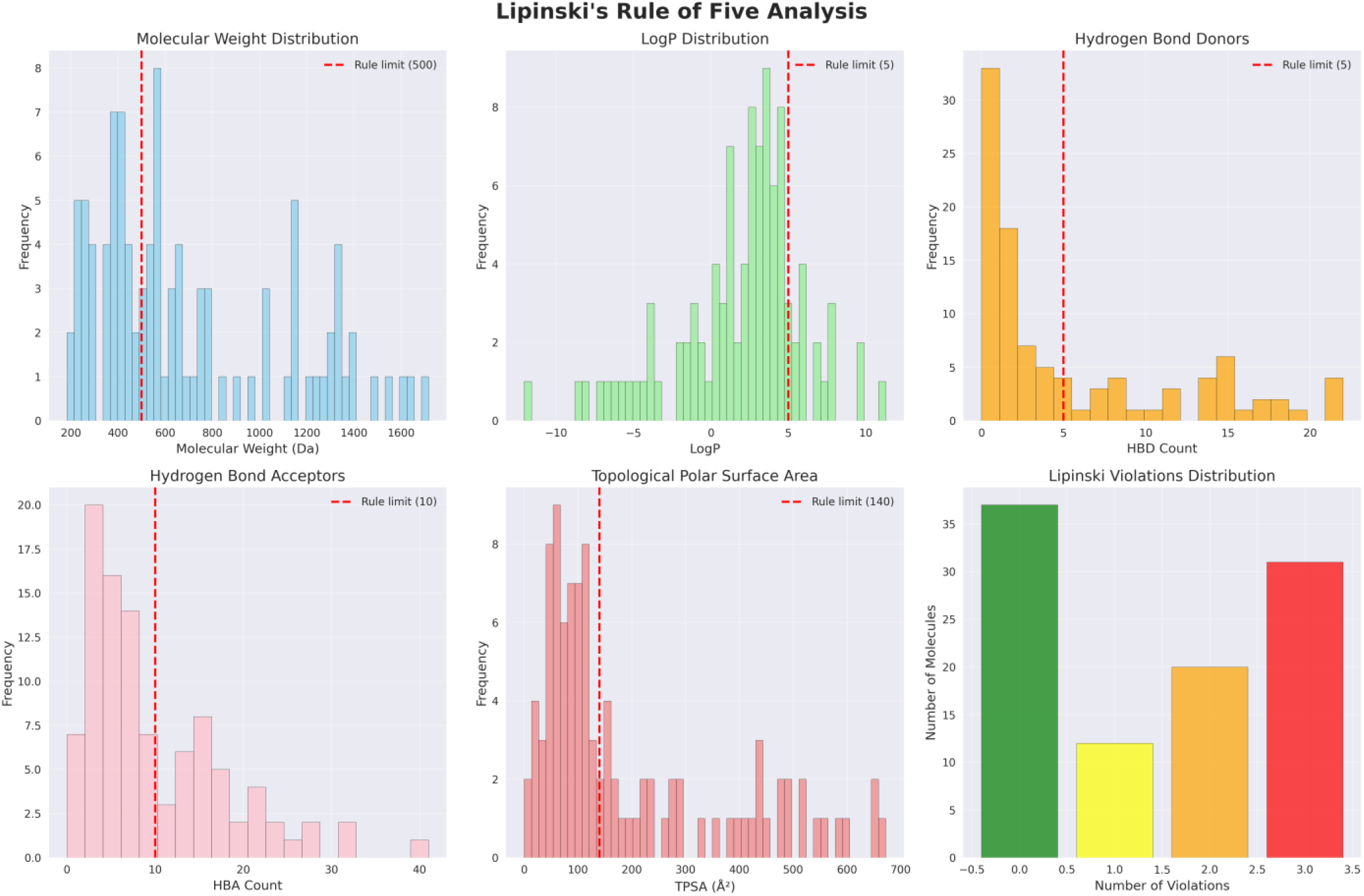
Lipinski Analysis of the ChEMBL database

### 3.2 Virtual Screening Results and Binding Affinity Landscape

High-throughput GPU-accelerated docking of 1,423,814 filtered compounds against the ThiE binding pocket revealed a binding affinity distribution with maximum and minimum affinities of -51.4 kcal/mol and 995.9 kcal/mol respectively (Figure 3). Top 10 resulting molecules were further subjected to lead optimization (Figure 4).

**Figure 3:**
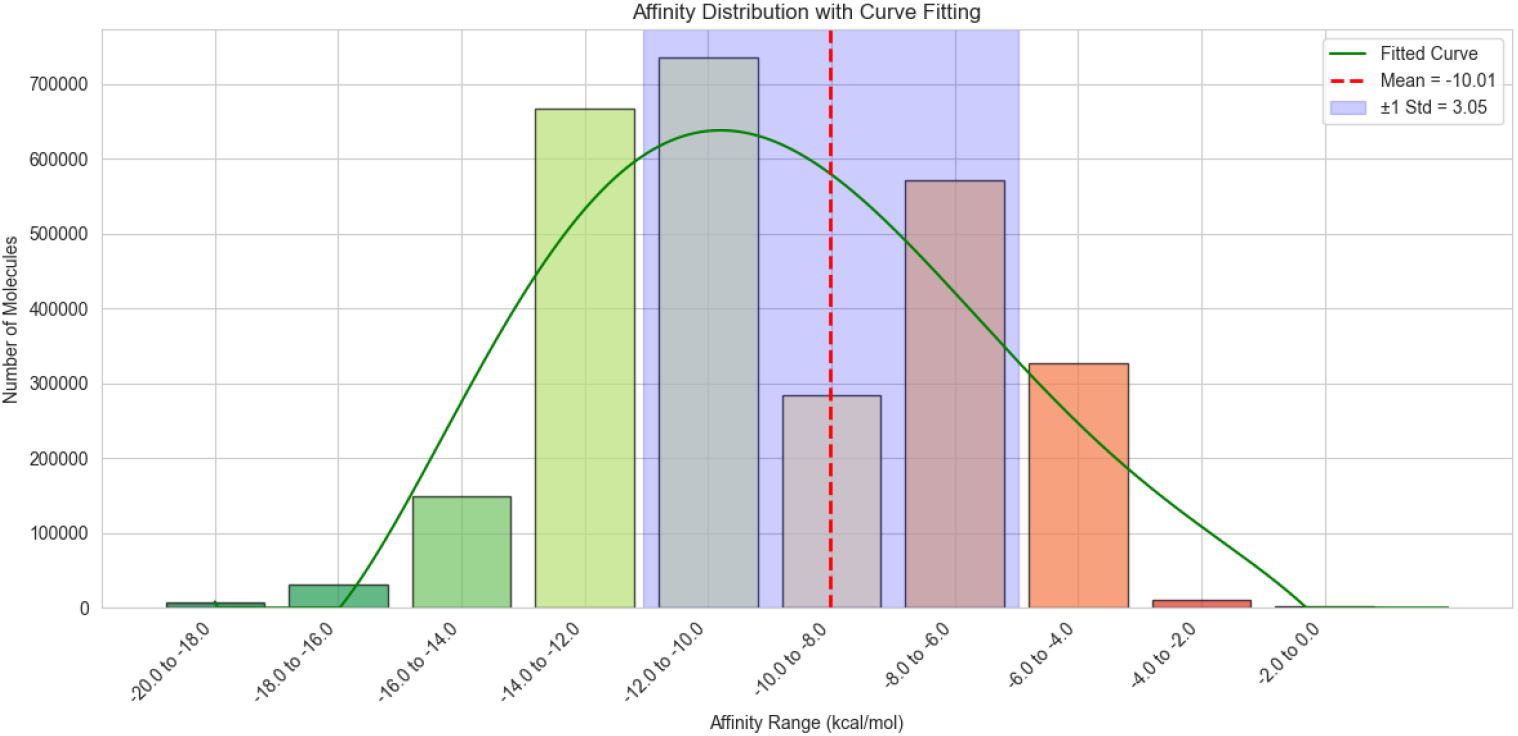
Binding Affinity Distribution

**Figure 4:**
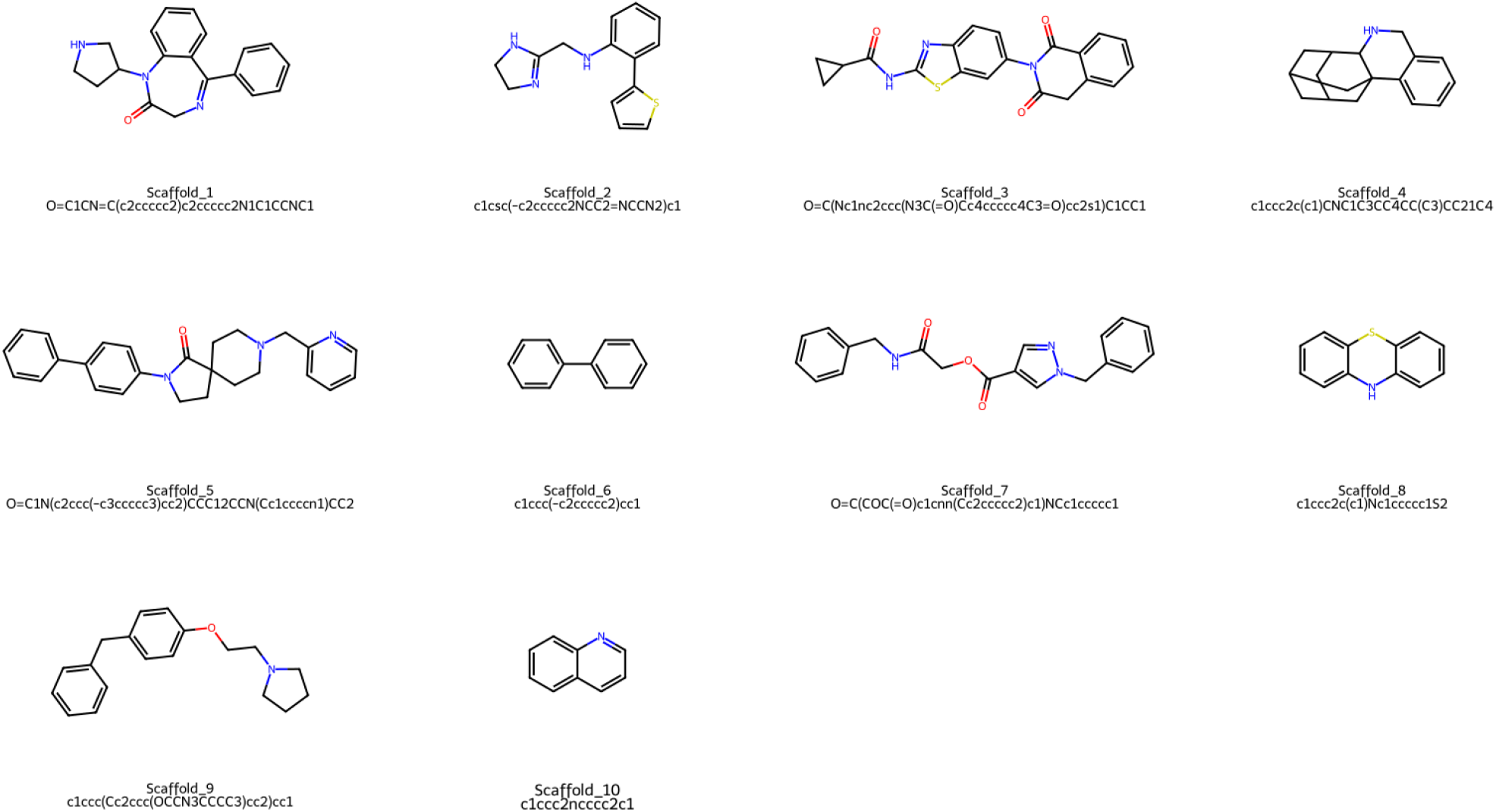
Top 10 molecules obtained by virtual screening sorted according to binding affinity

### 3.3 Structure-Activity Relationship Analysis: Scaffold Identification and R-Group Effects

R-group substitution of R1 and R2 of 10 scaffolds with 10 different functional groups provided 1,000 unique molecular combinations (Supplemental Figure 2). IC_50_ prediction was performed for each combination using Boltz-2 affinity module. The resulted affinity displayed a mean log_10_IC_50_ of 1.027 in µM and a lowest value of -0.201 corresponding to predicted IC50 values of 10.64 µM and 630 nM respectively (Figure 5).

**Figure 5:**
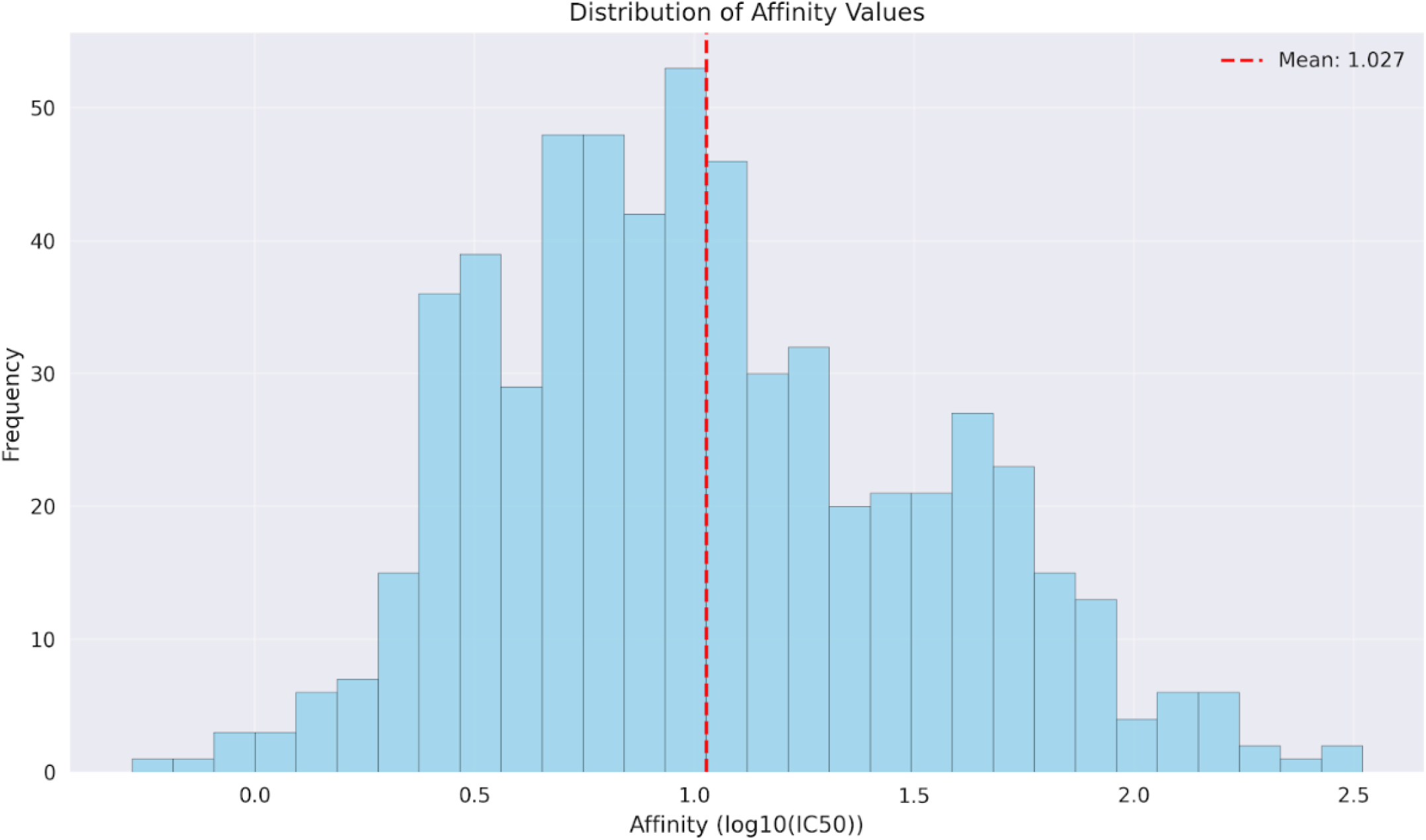
Predicted IC_50_ distribution of molecular mutants

Analysis of R-group substitution effects revealed clear patterns in how substituent functional groups modulate binding affinity. For example, within Scaffold 9, R1 and R2 substitution effects demonstrate that a dual combination of either strongly aromatic hydrophobic or negatively charged hydrophilic substituents have significantly improved affinity. Conversely, complementary combinations of those substituents too show modest affinity enhancements. Meanwhile, in Scaffold 8, amphiphatic combinations show significant affinity gains, but dual combinations of similar substituents show significant limitations (Figure 6).

**Figure 6:**
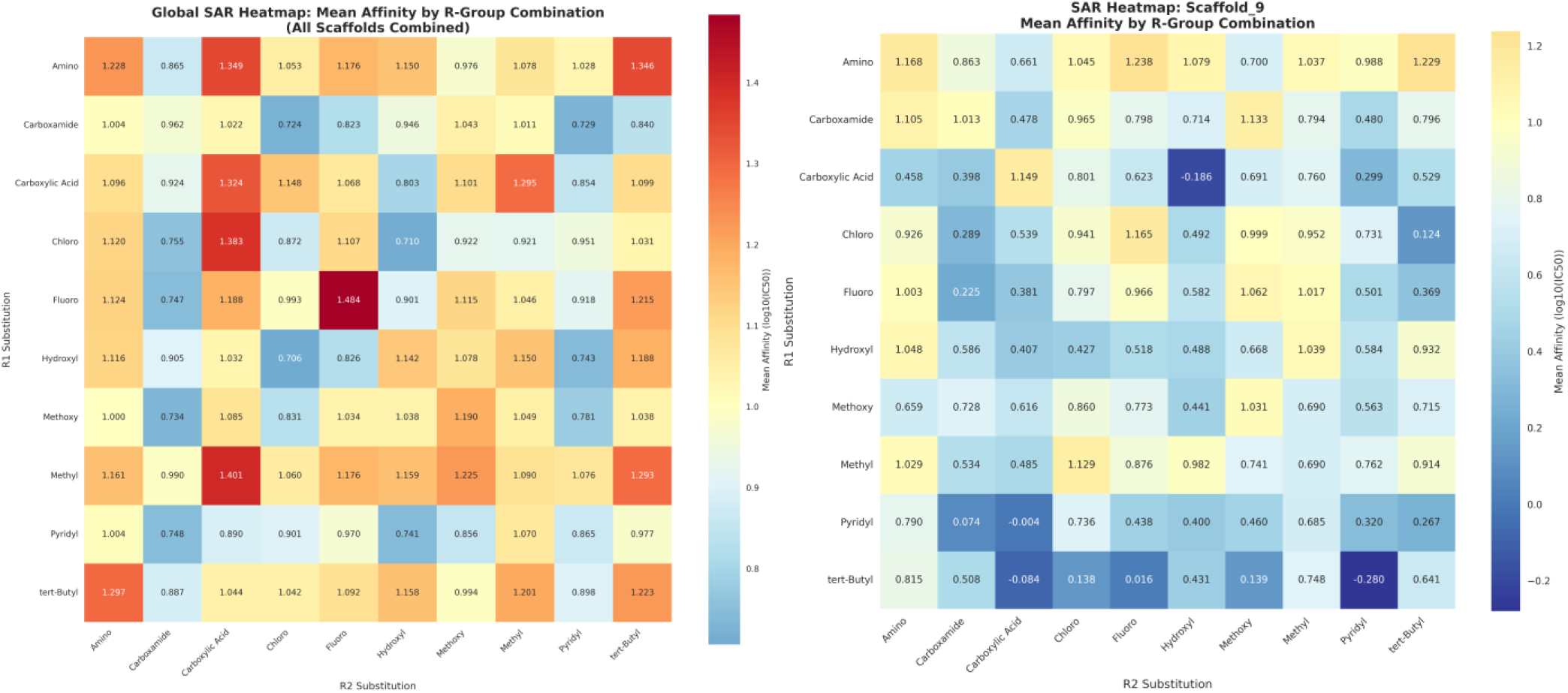
SAR Analysis of mean of 10 Scaffolds and Scaffold 9.

The non-additive nature of R1 and R2 combination effects, substantially exceeding the predicted additive combination of individual substituents, imply possible effects of intramolecular interactions within the substituents upon the overall binding affinity (Figure 7).

**Figure 7:**
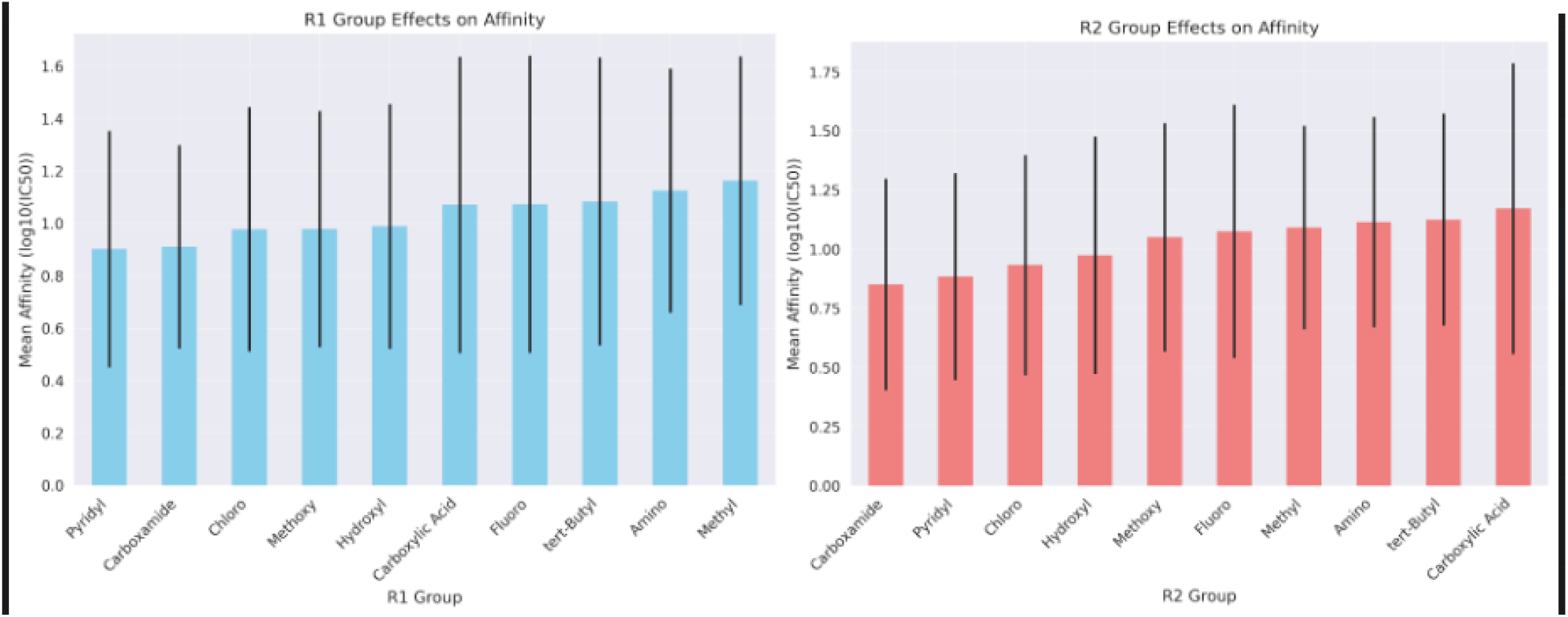
Effect of each R-Group on predicted IC_50_ across 10 scaffolds at R1, R2

Figure 8 displays the top 20 molecules sorted according to predicted log_10_IC_50_.

**Figure 8:**
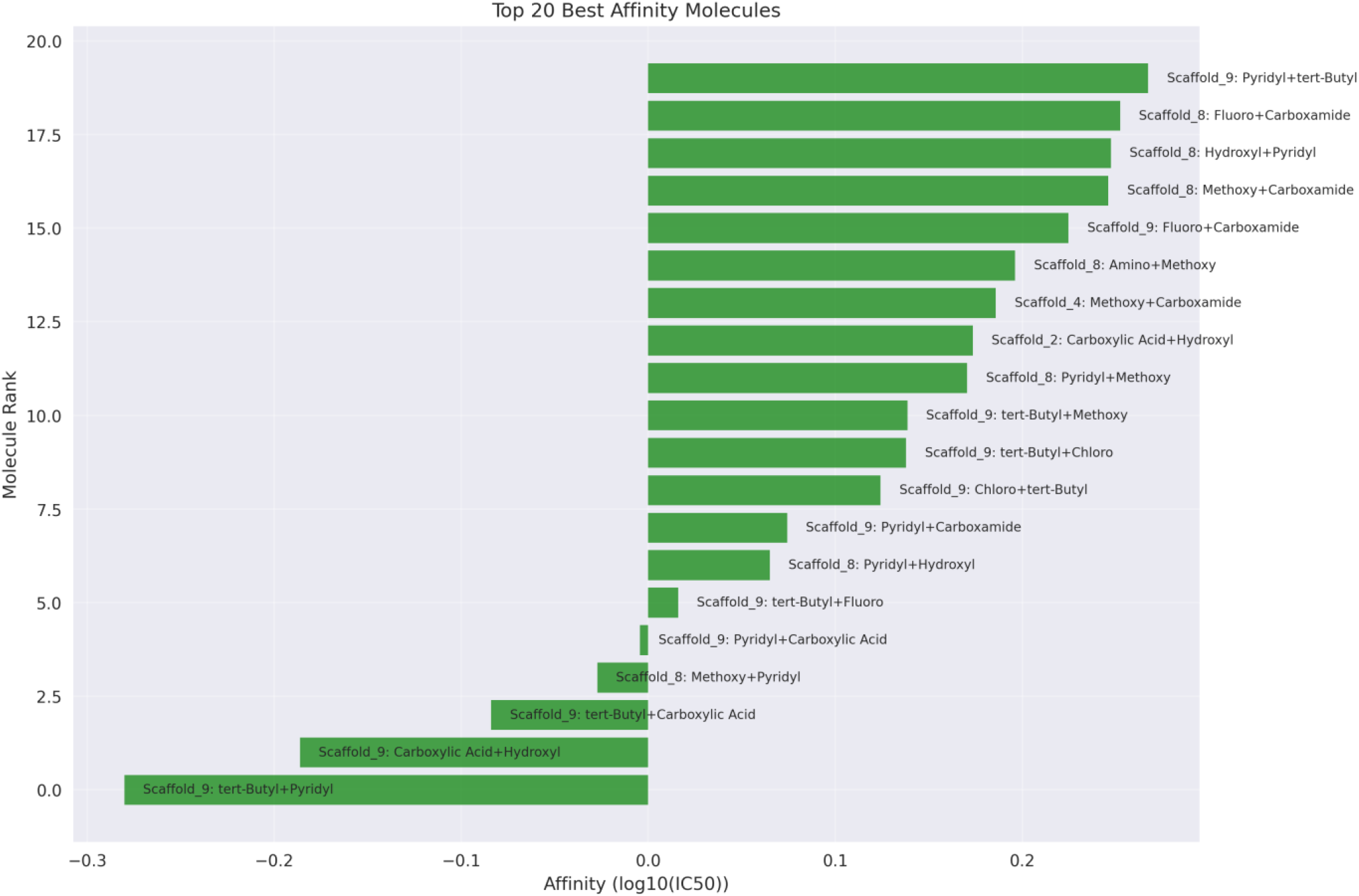
Top 20 molecular mutants ranked according to predicted IC_50_

### 3.4 ADMET Profiling and Multi-Criteria Lead Selection

Application of the TOPSIS multi-criteria decision analysis identified the molecule code-named NK1 (Figure 9) as the leading candidate with the highest TOPSIS score among them.

**Figure 9:**
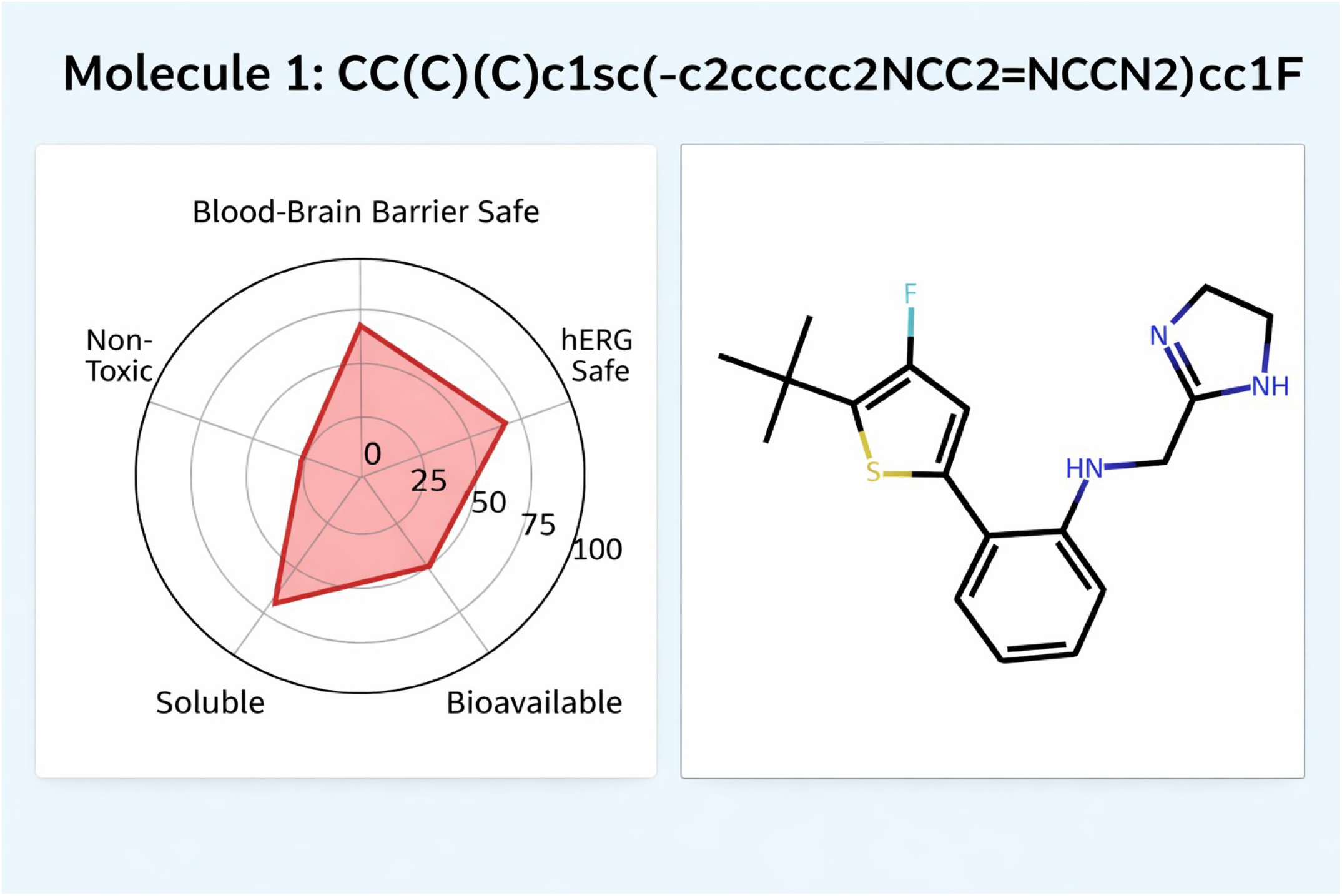
NK1 obtained with highest TOPSIS score

Representative ADMET properties of NK1 were LogP = 3.032, predicted aqueous solubility (logS) = 3.791 (indicating favorable solubility in physiological media), Synthetic Accessibility score = “Easy” (indicating synthetic tractability for experimental validation), Caco-2 permeability predicted as favorable (25 nm/s), suggesting good intestinal epithelial absorption, BBB penetration score predicted as “Safe” with low probability of central nervous system penetration, minimizing neurotoxicity risk, hERG blocker prediction indicating no blocking activity, reducing cardiac toxicity potential, hepatotoxicity prediction indicating “Non-Toxic” with low probability of liver injury, AMES mutagenicity prediction indicating “Non-Toxic”.

### 2.5 Molecular Dynamics Simulation Results and Binding Stability

Key stability metrics from the molecular dynamics simulation of the protein-ligand complex of NK1 over 100 nanoseconds were:

Mean RMSD of the holoprotein was 1.48 Å (Supplemental Figure 3), implying relatively low structural deviation from the initial conformation. RMSD values rapidly stabilized within 30-40 ns of the production simulation. For comparison, apoprotein RMSD was found to be 1.38 Å (Supplemental Figure 4), suggesting that ligand binding has further stabilized the protein structure and reduced the total energy of the system, suggesting strong affinity towards the target protein.

Ligand all-atom RMSD was found to be 1.84 Å (Figure 10), indicating that the small molecule remains stably bound within the active site without undergoing spontaneous dissociation.

**Figure 10:**
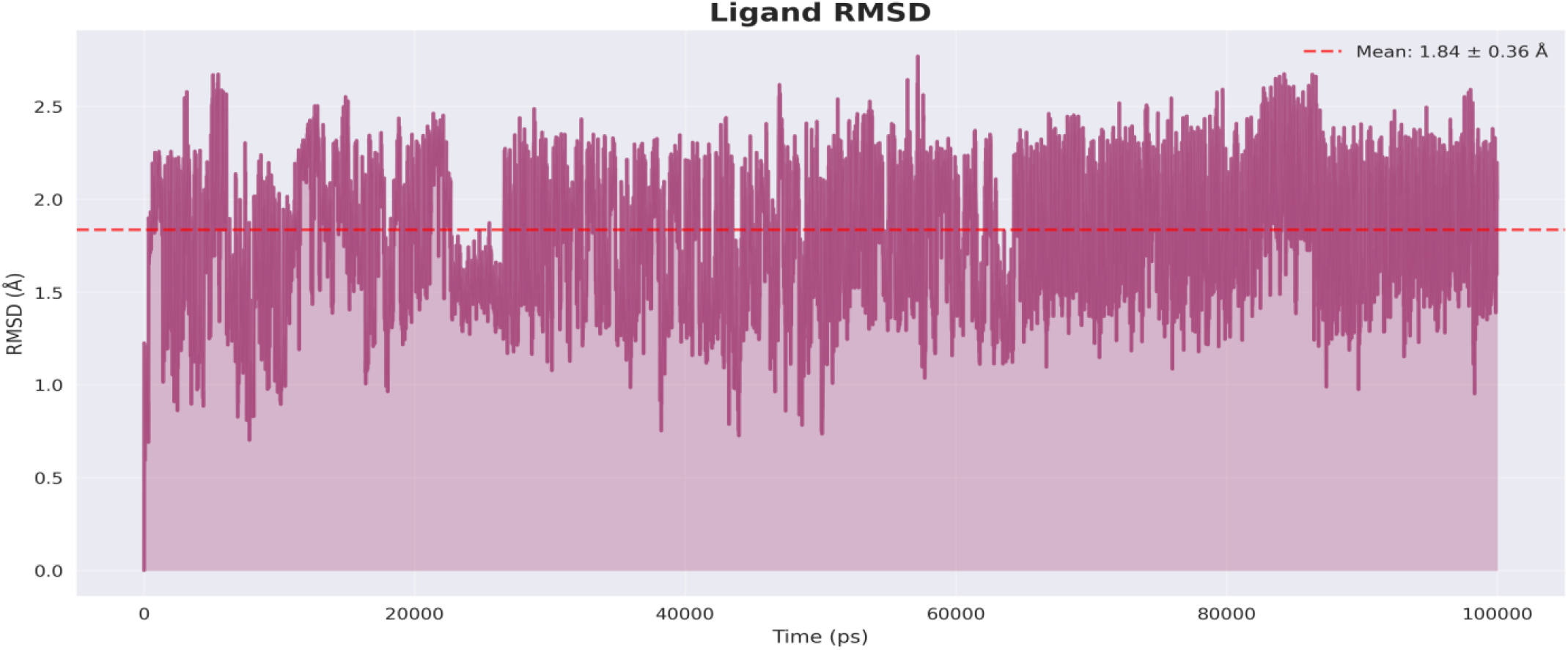
Ligand All-Atom RMSD

Radius of gyration of the protein remains near constant at 16.5 Å across the trajectory with negligible deviation (Supplemental Figure 5). Mean RMSF across all residues were 17.6 Å calculated from the center of mass of the protein with distinct regional variation (Supplemental Figure 6). Residues part of loops and terminal regions were highly flexible whereas the conserved catalytic core residues (N69, D70, D89, S108, Q135, K137, G166, V186) were highly constrained. Particularly, the residues directly contacting the bound inhibitor exhibited very low RMSF values.

## 4. CONCLUSIONS

This present investigation extends and complements prior research establishing thiamine biosynthetic enzymes as antimicrobial targets. Our computational identification of multiple high-affinity scaffolds with K_D_ predictions in low micromolar range opens avenues for exploring thiamine biosynthesis pathway as a viable drug target for future generation antimicrobials. The observation that ThiE orthologs from diverse bacterial species display high structural conservation of backbone RMSD 1.6 Å despite 30% sequence identity suggests that optimized inhibitors identified in this study may possess broad-spectrum efficacy. High-throughput virtual screening of more than 1.5 million molecules provides comprehensive chemical diversity coverage. During lead optimization, the systemic observation that aromatic pyridyl substituents consistently enhance binding affinity compared to alternatives. This suggests that pi-pi stacking and particularly with the histidine residues within the binding pocket largely contributes to the binding energy. Within Scaffold 9, both dual combination of similar substituents and amphiphatic combinations significantly enhance affinity. This indicates that the amphiphatic nature of the site of growth, with both entropy-driven (hydrophobic groups forming pi-pi interactions and Van der Waals interactions) and enthalpy-driven (hydrophilic groups forming strong hydrogen bonds and salt bridges) energies contributing to the overall affinity. This provides crucial information regarding engineering the ligand heavy atoms interacting with this region to maximize affinity. The molecular dynamics results specifically confirm that the binding conformation remains stable under realistic physiological conditions without spontaneous dissociation over 100 ns. Further experimental in vitro assays must be performed to assess the clinical potential of the molecules.

## ACKNOWLEDGEMENTS

The authors acknowledge SASTRA Deemed to be University for the infrastructural support and the computational resource facility - TIFAC Core.

