## Supplemental File for "Structure-Based Drug Design of Novel Inhibitors Targeting the Thiamine Biosynthesis Pathway in Bacteria"

### Supplemental material

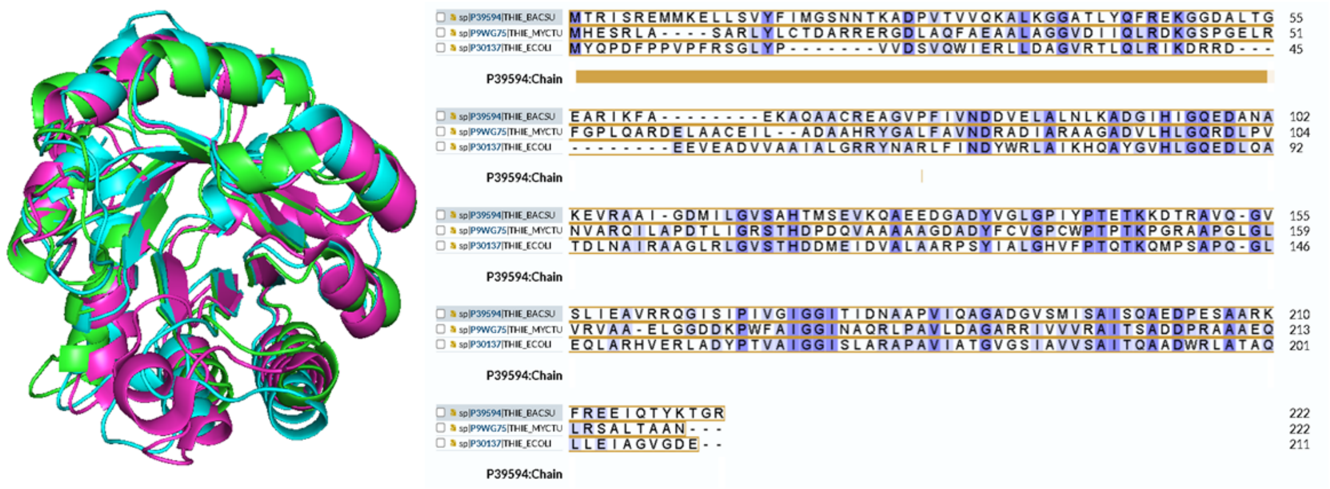


**Supplemental Figure 1:** Structural Superposition and Multiple Sequence aligment of ThiE proteins from *E. coli* (green)*, M. tuberculosis* (pink)*, B. subtillis* (blue)

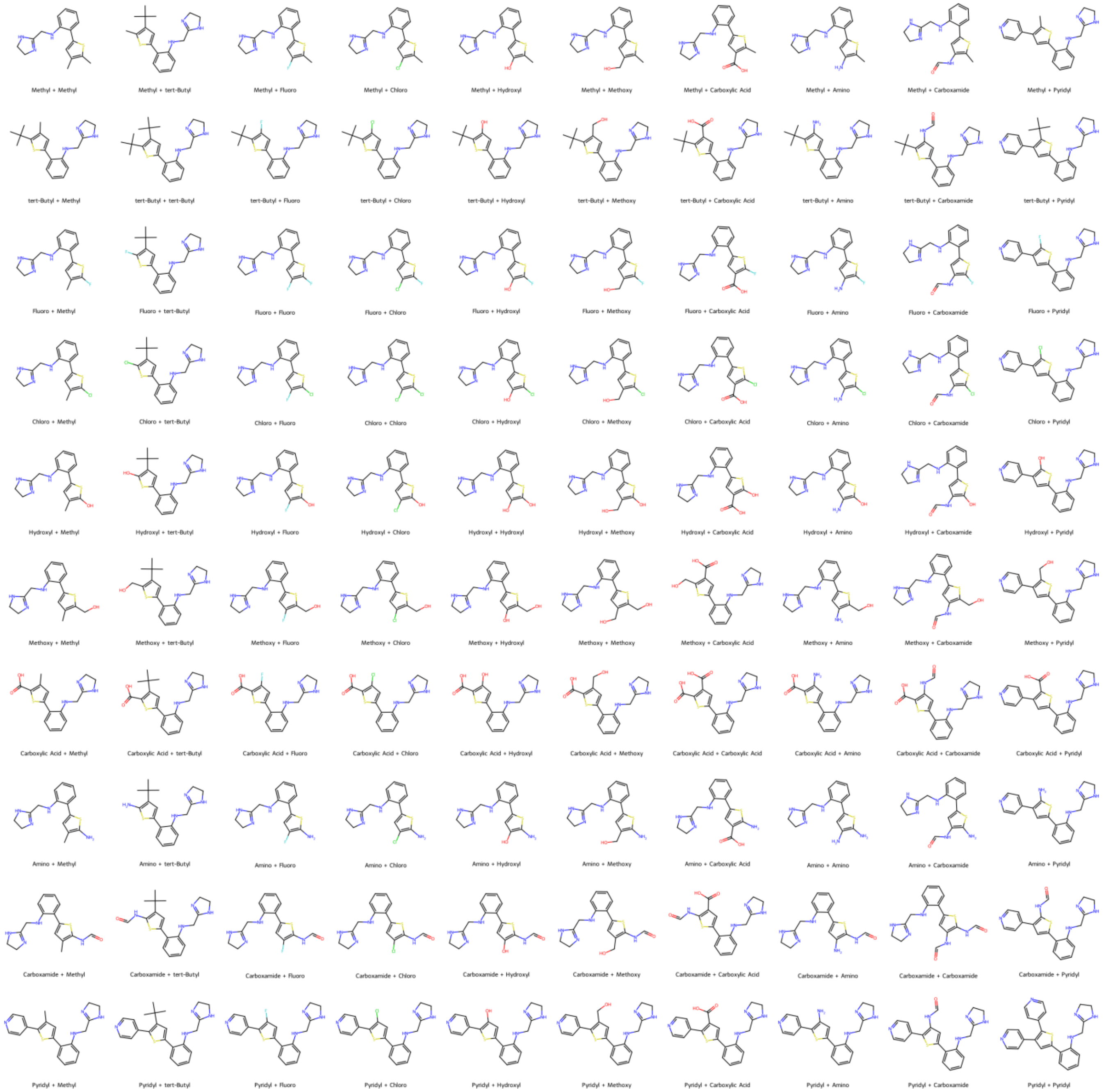


**Supplemental Figure 2:** Molecular Mutants produced by R-group substitution at R1, R2 in Scaffold 2

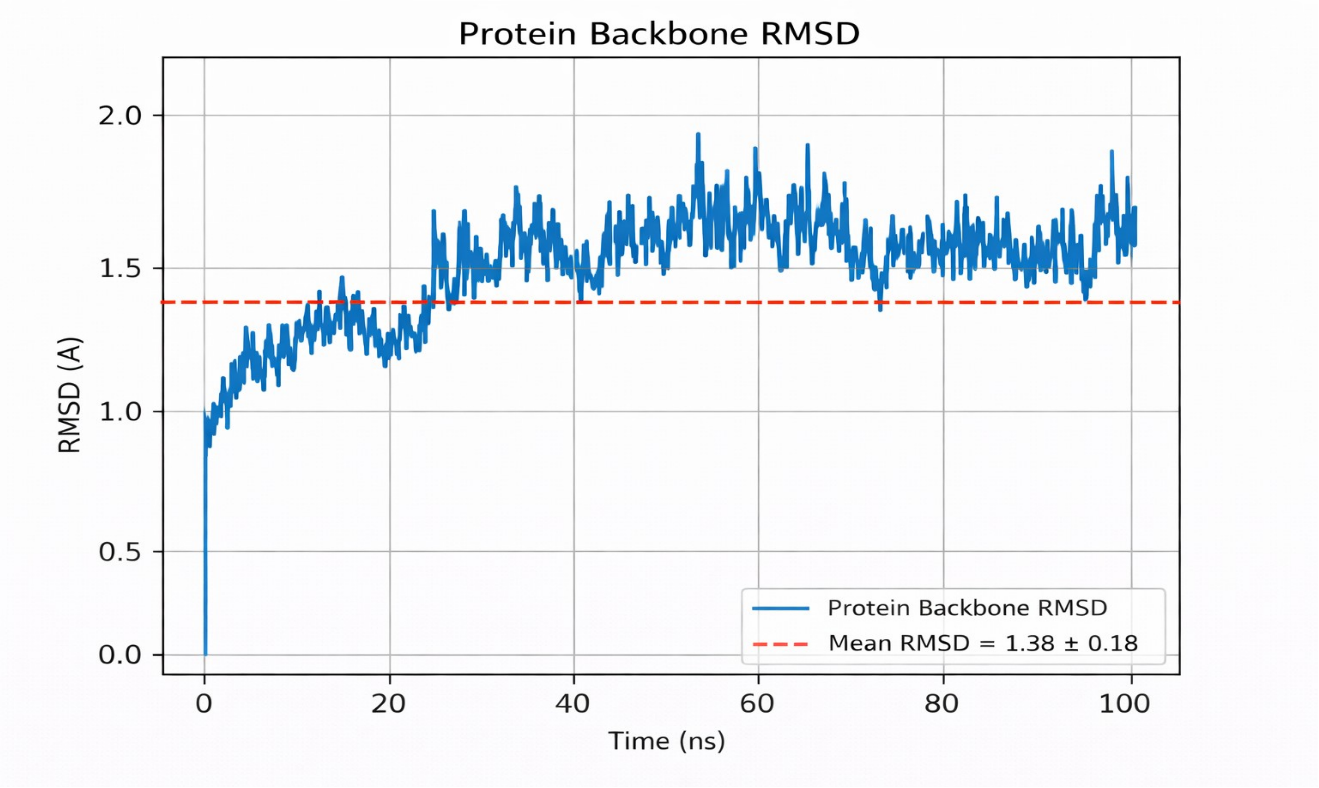


**Supplemental Figure 3:** Holo Protein Backbone RMSD

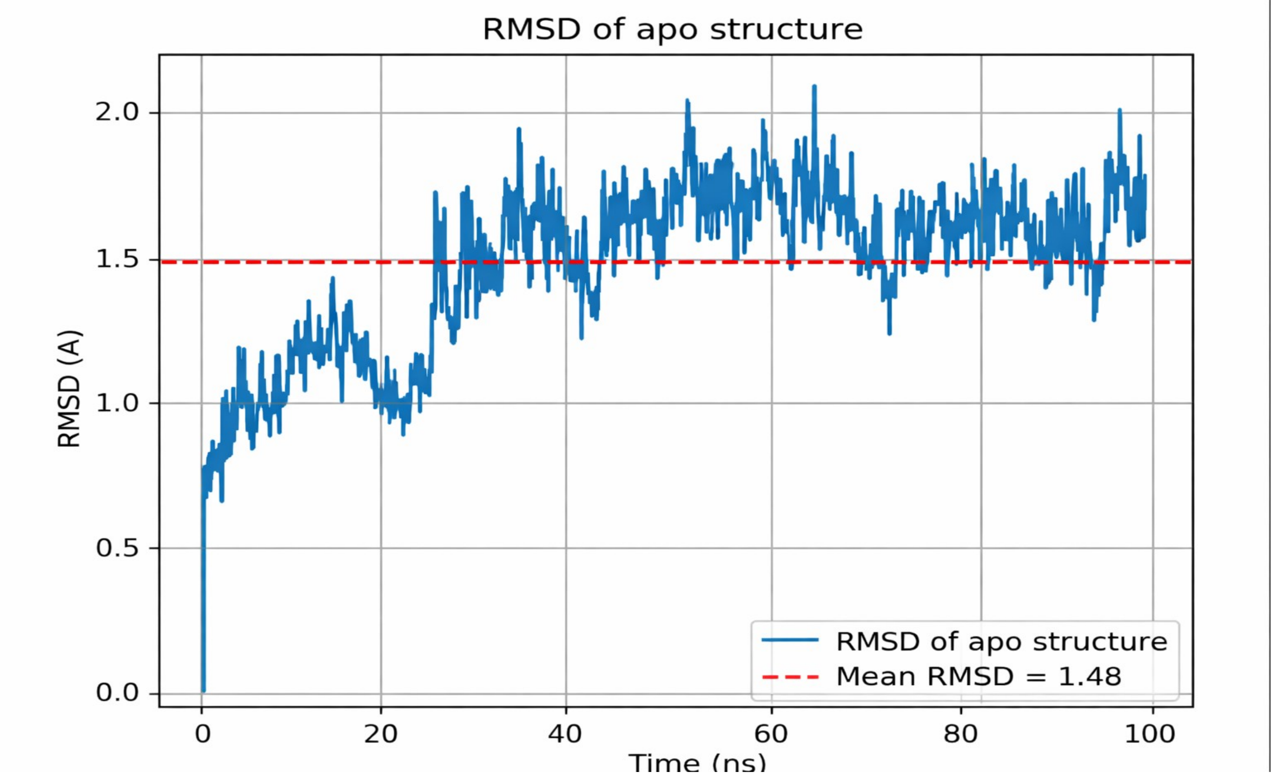


**Supplemental Figure 4:** Apo Protein Backbone RMSD

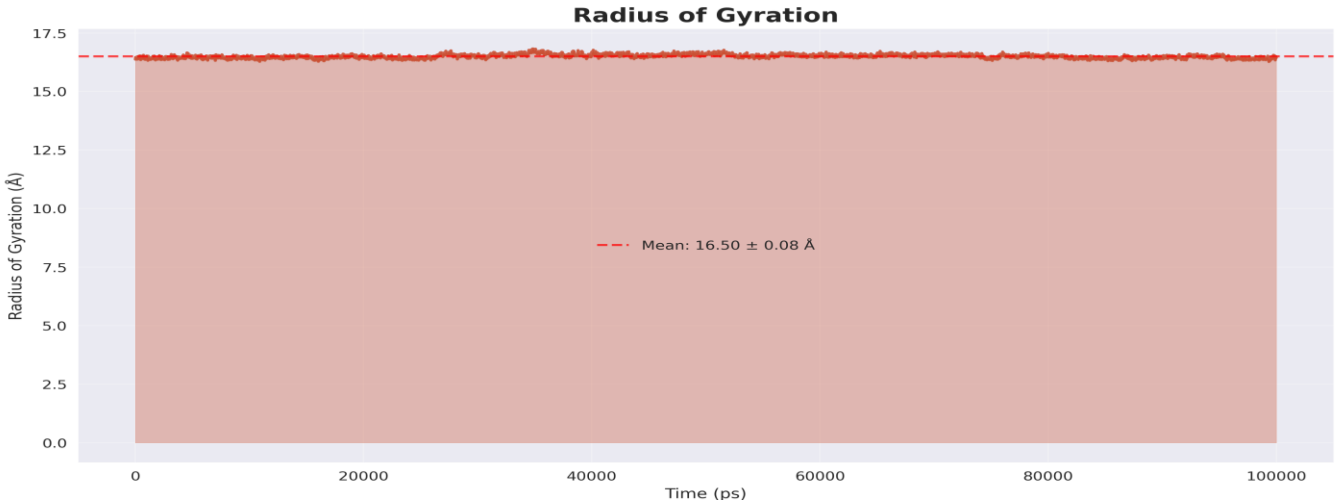


**Supplemental Figure 5:** Radius of Gyration

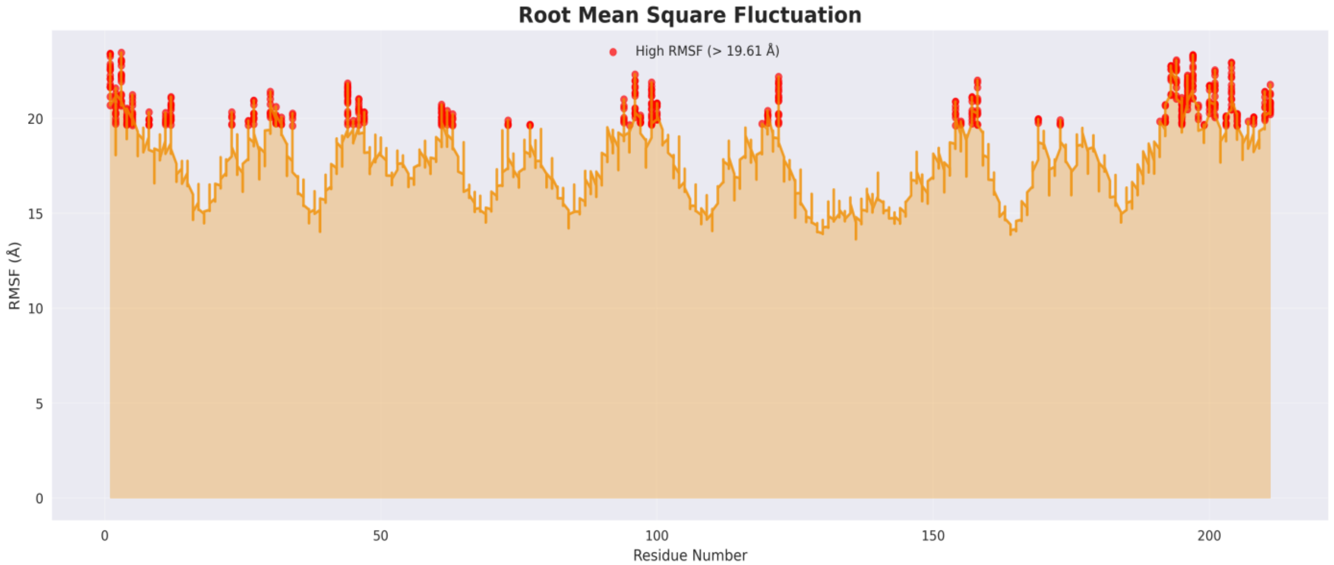


**Supplemental Figure 6:** Protein residue RMSF
